# Paternal-age-related de novo mutations and risk for five disorders

**DOI:** 10.1101/192740

**Authors:** Jacob L. Taylor, Jean-Christophe P.G. Debost, Sarah U. Morton, Emilie M. Wigdor, Henrike O. Heyne, Dennis Lal, Daniel P. Howrigan, Alex Bloemendal, Janne T. Larsen, Jack A. Kosmicki, Daniel J. Weiner, Pediatric Cardiac Genomics Consortium, Jason Homsy, Jonathan G. Seidman, Christine E. Seidman, Esben Agerbo, John J. McGrath, Preben Bo Mortensen, Liselotte Petersen, Mark J. Daly, Elise B. Robinson

## Abstract

**Background:** There are well-established epidemiologic associations between advanced paternal age and increased offspring risk for several psychiatric and developmental disorders. These associations are commonly attributed to age-related *de novo* mutations. However, the actual magnitude of risk conferred by age-related *de novo* mutations in the male germline is unknown. Quantifying this risk would clarify the clinical and public health significance of delayed paternity.

**Methods:** Using results from large, parent-child trio whole-exome-sequencing studies, we estimated the relationship between paternal-age-related *de novo* single nucleotide variants (dnSNVs) and offspring risk for five disorders: autism spectrum disorders (ASD), congenital heart disease (CHD), neurodevelopmental disorders with epilepsy (EPI), intellectual disability (ID), and schizophrenia (SCZ). Using Danish national registry data, we then investigated the degree to which the epidemiologic association between each disorder and advanced paternal age was consistent with the estimated role of *de novo* mutations.

**Results:** Incidence rate ratios comparing dnSNV-based risk to offspring of 45 versus 25-year-old fathers ranged from 1.05 (95% confidence interval 1.01–1.13) for SCZ to 1.29 (95% CI 1.13-1.68) for ID. Epidemiologic estimates of paternal age risk for CHD, ID and EPI were consistent with the dnSNV effect. However, epidemiologic effects for ASDs and SCZ significantly exceeded the risk that could be explained by dnSNVs alone (p<2e-4 for both comparisons).

**Conclusion:** Increasing dnSNVs due to advanced paternal age confer a small amount of offspring risk for psychiatric and developmental disorders. For ASD and SCZ, epidemiologic associations with delayed paternity largely reflect factors that cannot be assumed to increase with age.

---

Epidemiologic studies consistently link advanced paternal age to risk for psychiatric and developmental disorders in the next generation. This effect is particularly well characterized in autism spectrum disorders (ASDs) and schizophrenia (SCZ), where the children of older fathers have been found to be at increased risk for both disorders in many large, population-based studies.^1-7^ Advanced paternal age has also been associated with risk for several other early-onset disorders, including congenital heart disease (CHD),^8^ epilepsy,^9^ and intellectual disability (ID).^10^

It is commonly suggested that these observations reflect the influence of *de novo* mutations—genetic variants that are new to a family and usually appear during the formation of germ (sperm and egg) cells.^2,11,12^ *De novo* genetic variants, especially *de novo* single nucleotide variants (dnSNVs), arise in paternal germ cells 3-4 times more often then they do in maternal germ cells.^11,13^ As men age, dnSNVs accumulate in their sperm.^14^

Paternal-age-related *de novo* variants are therefore likely to increase risk for any dnSNV-influenced disorder. However, the size of that risk is unclear, and could in fact be far smaller than that suggested by the epidemiologic associations. This possibility received support in a recent study using simulated data that found that common polygenic risk, rather than *de novo* variation, could be most relevant to the epidemiologic patterns observed for ASDs and SCZ.^15^ This finding is consistent with the hypothesis that common, inherited genetic risk for psychiatric disorders may also predict age of childbearing.^16,17^ That is, individuals who carry elevated, *inherited* risk for ASDs, for example, might on average have children later in life.^1^

It is now possible to approach this problem using empirical data, as, over the last several years, large parent-child trio whole-exome-sequencing studies have been able to quantify the relationship between *de novo* variation and risk for each of ASDs, CHD, ID, neurodevelopmental disorders with epilepsy (EPI) and SCZ.^18-33^ The purpose of this study was to directly estimate risk for each of these five disorders created by paternal-age-related *de novo* mutations in the exome, the protein-coding portion of the genome. Clarity regarding the *amount* of *de novo* mutation risk conferred by advancing paternal age could influence individual decision-making and genetic counseling.^34^

We next conducted an analysis in the population-based Danish national patient registry to comparably estimate the epidemiologic association between advanced paternal age and risk for each disorder. In so doing, we noted substantial variation in the fraction of the epidemiologic effect that could be explained by *de novo* mutations in the exome. We found that the epidemiologic associations for CHD, EPI and ID were consistent with, and could plausibly be attributed to, the effects of accumulating dnSNVs. However, the associations seen for disorders that specifically impact social behavior – ASDs and SCZ – significantly exceeded the amount of risk that could be explained by *de novo* mutations.

## Methods

### Estimating risk from paternal age related *de novo* mutations in the exome

We developed a statistical model that, for fathers of any two given ages, compares offspring disease risk due to nonsynonymous (missense and protein truncating) dnSNVs. For the remainder of the text, “dnSNV” will be used to reference nonsynonymous dnSNVs, as they are the *de novo* variant types consistently associated with risk for human disease.^18-33,35^ The model incorporates two relationships: 1) the relationship between paternal age and dnSNV accumulation and 2) the relationship between dnSNVs and the outcome of interest (e.g. ASDs).

We first empirically estimated the general population relationship between paternal age and number of dnSNVs in offspring. The analysis used data from trio-sequenced healthy siblings of ASD probands in the Simons Simplex Collection (SSC), which has been described extensively.^18,36^ To our knowledge, the SSC siblings constitute the largest existing sample of control individuals with available whole-exome dnSNV data.

Using poisson regression, we estimated the association between number of offspring per year of increasing paternal age (Box 1; Figure S1a). We did not include maternal age as a covariate in the model, as paternal and maternal ages were too highly correlated (r = 0.72) to distinguish between their effects (Supplementary Appendix A), and the SSC *de novo* variant data have not been phased in adequate fraction for the effects to be estimated separately.^18^ While the maternal age effect on dnSNV accumulation is much smaller than the paternal age effect,^11,13^ this modeling approach results in estimates of paternal-age risk that include (and are increased by) the correlated risks associated with maternal age (Supplementary Appendix A). As the model is designed to predict changes in disease incidence at a population-level, the results presented here will best generalize to populations in which maternal and paternal age have a similar relationship to that observed in the United States.

We next estimated the relationship between dnSNVs and offspring risk for each of ASD, CHD, EPI, ID, and SCZ – the disorders for which large-scale whole-exome-sequenced trio data currently exist. All such data has been published^18-32^ with the exception of dnSNV data from 1135 out of 2348 CHD trio families. Data from these families have been collected through the Pediatric Cardiac Genomics Consortium (PCGC) (Supplementary Appendix B). From each study, or group of studies, we extracted the average number of dnSNVs per case. To estimate an average dnSNV rate in controls, we again used the healthy siblings of probands from the SSC.

The statistical model is presented in Box 1, and is described in detail in Supplementary Appendix B. The output of the model is an incidence rate ratio (IRR) reflecting the increase in dnSNV-related disease risk in offspring of older fathers compared with offspring of younger fathers.

#### Box 1 Predicting changes in population-level disease incidence due to paternal age related dnSNVs in the exome

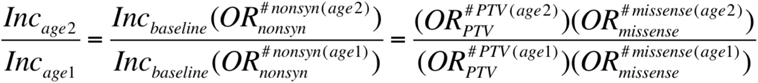

Where:

*#nonsyn(age)* = number of dnSNVs expected in offspring of fathers of a particular age *=e*^*i+age*β*^ (see Supplementary Appendix B). *OR_nonsyn_ / OR_PTV_ / OR_missense_* = odds ratio reflecting disease risk associated with having one additional nonsynonymous, protein truncating or missense variant respectively

In brief, one can estimate disease incidence in offspring of fathers of a specified age (e.g. people born when their fathers were 25) by multiplying (a) disease incidence among individuals who carry zero dnSNVs (INC_baseline_) and (b) the odds ratio (OR) associated with one additional nonsynonymous dnSNV, exponentiated to the expected number of dnSNVs in offspring of fathers of the specified age. Part (b), presented in parentheticals in Box 1, is estimated empirically through the inputs described above. Baseline incidence can be unknown as, algebraically, it is not needed for estimating the IRR. As further noted in Box 1, effect size variation across different types of dnSNVs is also algebraically irrelevant since they all increase with age at the same rate. To show this we use the example of missense and protein truncating variants (PTVs), which have different average effect sizes on risk for ASD. The principle is relevant to all potential subclasses of dnSNVs, as long as each subclass increases at the same rate with respect to paternal age (Supplementary Appendix C). As described in supplementary Appendices D and E, the model’s estimates are robust to: plausible variation in the expected (control) rate of dnSNVs, plausible variation in the estimated effect size of case dnSNVs, and the inclusion of *de novo* copy number variants. The model assumes that risk-conferring statistical interaction among dnSNVs do not commonly occur. In supplementary Appendix F we show that such interactions are not currently identifiable in the ASD data and discuss why they are unlikely to be relevant for any of the other disorders.

We used the model to specifically estimate the IRR for each disorder in offspring of 35-year old, 45-year old and 55-year old fathers compared with offspring of 25-year old fathers. 95% confidence intervals were determined by simulating 10,000 versions of each input parameter using the actual estimated parameters and associated standard errors (Supplementary Appendix G). As further described in supplementary Appendix H, the confidence intervals illustrate the range of possibilities for each comparison (e.g. offspring ASD risk between fathers aged 45 v. 25) but are not valid for comparing estimates of the dnSNV effect either within or across disorders.

### Estimating the epidemiological associations between advanced paternal age and disorder risk using the Danish national registries

We used the Danish national psychiatric and patient registries to estimate a comparable epidemiologic association between increasing paternal age and risk for each of the five disorders. Previous reports have used the Danish registries to estimate paternal age effects for ASDs, schizophrenia and CHD.^1,37^ Our approach is similar to these earlier efforts, but modified slightly to maximize concordance with our dnSNV model. We queried the registries for all individuals born in Denmark to Danish-born parents between 1955 and 2012 with a diagnosis of ASD, EPI, ID, or SCZ issued between 1995 and 2012, such that all diagnoses were made using ICD-10 criteria. Individuals with any of the above diagnoses made prior to 1995 were excluded. To be consistent with the PCGC exome sequencing analysis, CHD cases were only included if diagnosed within one year of birth, and were therefore born between1994-2011. The controls for the CHD analysis were also all born between 1994-2011. For all five disorders, we aimed to identify ICD-10 diagnoses that were as similar as possible to the phenotypes included in the exome sequencing studies listed in Table 1 (Supplementary Appendix I and Tables S4 and S5). For each of the five disorders, all non-cases born within the time period were used as controls (n > 300,000 for each analysis).

To permit a comparison between the epidemiologic data and our dnSNV model, we identified all case and control individuals whose fathers’ age at the time of their birth was either 20-29 or >39. The broad paternal age bins were designed to ensure a sufficient number of cases of each disorder per bin for statistical comparison. We set a lower age bound of 20 because very young parent age is also associated with risk for several of the five disorders, though likely through different mechanisms.^1,10^

We used Cox proportional hazard models to compare offspring risk for ASD, EPI, ID, and SCZ between fathers in the two different age bins. Age was used as the primary time scale. As in the dnSNV model, we did not control for maternal age in the primary analyses. We repeated the analysis controlling for maternal age and/or calendar period and neither produced a substantial difference in our estimates (Supplementary Appendix J and Table S6). Since age of diagnosis was constrained for CHD, we estimated ORs across the paternal age groupings, rather than using a Cox proportional hazard model. HRs are functionally equivalent to IRRs,^38^ and the same is true for ORs for disorders with low cumulative incidence (e.g. <10%) and risk factors with small effect sizes.^39^ From that, we are able to meaningfully compare the epidemiologic estimates from the Danish data to IRRs estimated through the dnSNV model.

To compare estimates from the dnSNV and Danish epidemiologic models, we input the mean paternal ages of the Danish fathers into the dnSNV model, and ran the model for each disorder. For ASD, EPI, ID and SCZ, the mean age for each bin was calculated among all fathers of children born between 1955-2012. For CHD, the mean age for each bin was calculated among fathers of children born between 1994-2011.

To test whether the epidemiologic and dnSNV estimates were significantly different, we used standard errors of the natural log of the HRs (or OR) from the epidemiologic model to generate 100,000 plausible values for the true association (Supplementary Appendix K). As above, we used the standard errors for each parameter in the dnSNV model to generate 100,000 plausible values for the IRR between fathers with ages equal to the means of the two relevant age bins from the Danish data. We then used these distributions to generate p-values, by identifying the empirical probability of the observed divergences between the dnSNV and epidemiologic estimates, if the estimates were in actuality equivalent.

## Results

Using n=1827 control trios in whom data on paternal age was available, we found that dnSNVs accumulate with advancing paternal age at a rate of 3.1% per year (p=2e-10). This exome-specific estimate is consistent with those produced by recent whole genome sequencing studies (Table S1), and did not vary by type of dnSNV examined (Figure S1b).

Table 1 summarizes each data set used to estimate the risk associated with one additional dnSNV for each disorder. For this analysis there were n=1902 controls. For each disorder, cases had an excess of dnSNVs compared with controls (p < 0.05).

**Table 1:** The burden of dnSNVs in five disorders.

| Disorder | References | N trios | dnSNVs per person | OR (p) |
| --- | --- | --- | --- | --- |
| Intellectual disability | <sup>26-30,33</sup> | 5264 | 1.16 | 1.78 ($<1e^{-20}$) |
| Neurodevelopmental disorders with epilepsy | <sup>31</sup> | 2151 | 0.88 | 1.35 ($9e^{-17}$) |
| Congenital heart disease | <sup>32</sup>, unpublished data (see Supplementary Appendix B) | 2348 | 0.86 | 1.32 ($2e^{-14}$) |
| Autism spectrum disorders | <sup>18</sup> | 2508 | 0.79 | 1.22 ($3e^{-8}$) |
| Schizophrenia | <sup>19-25</sup> | 1077 | 0.72 | 1.11 (0.02) |
| Control | <sup>18</sup> | 1902 | 0.65 | - |

### Risk from paternal-age-related dnSNVs

Figure 1 illustrates the impact of paternal-age-related dnSNVs on offspring risk for each of the five disorders. For offspring of a 45 year old father compared with offspring of a 25 year old father, the IRR of each disorder ranged from: 1.05 (1.01 – 1.13) for SCZ to 1.29 (1.13-1.68) for ID (Table S2). This translates to an approximately 5% to 30% increase in risk over that paternal age span. This increase in risk must be interpreted against the low overall incidences of the five disorders.^30,31,40-42^ For example, in a condition with 1% baseline incidence, a 30% increase in risk would translate to a 1.3% probability of the outcome. The estimates from the model described here reflect overall risk for developing a disorder whether or not the disorder is caused by a *de novo* variant. If instead, we were to attempt to measure the increase in risk of developing each disorder only in circumstances where the disorder is *caused* by a dnSNV our estimates of the paternal age effect would be slightly higher, as they were in a recent analysis of the parental age effect on ID.^30^

**Figure 1:**
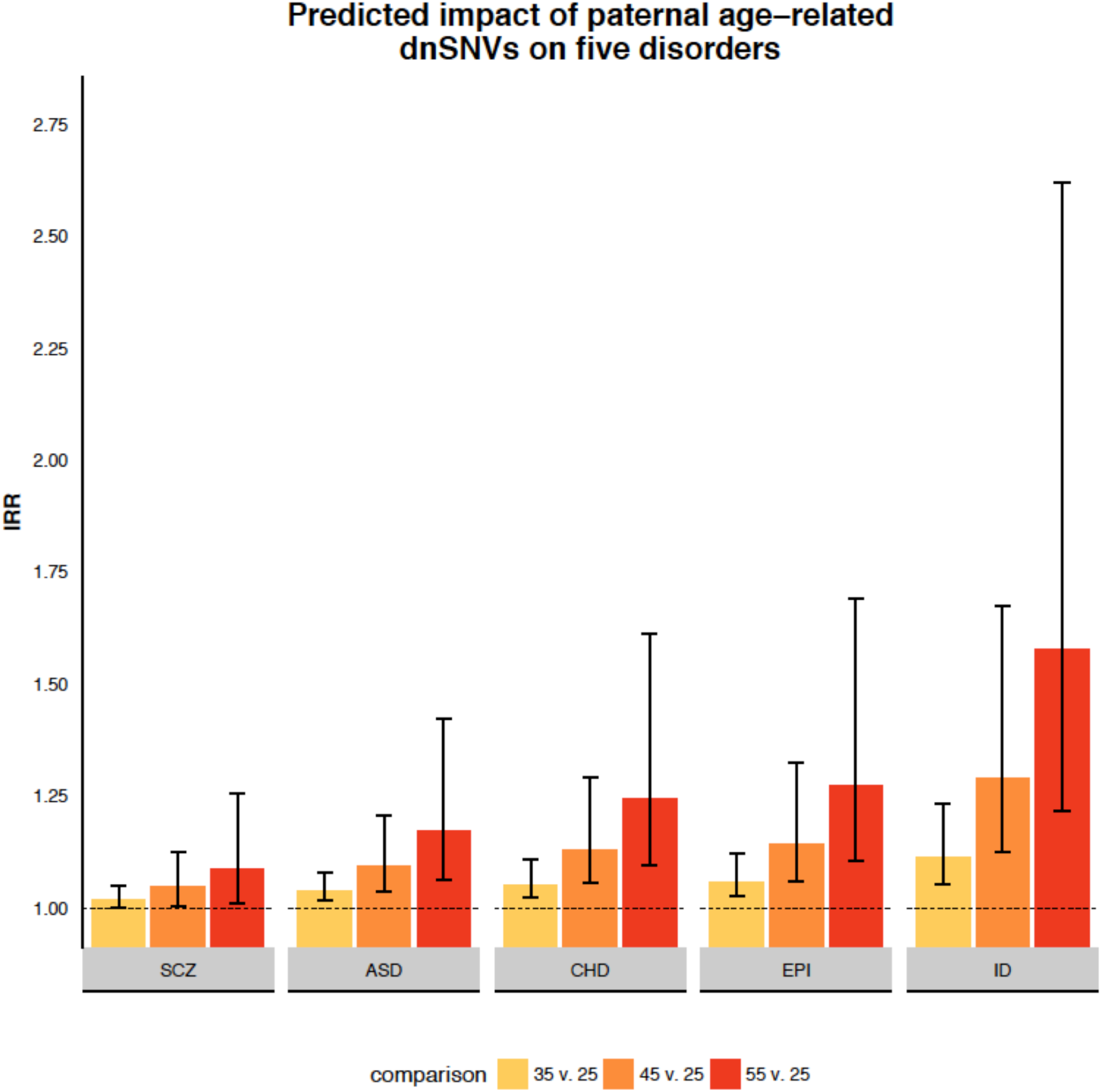
Note: IRR= Incidence rate ratio; SCZ = schizophrenia; ASD = autism spectrum disorder; CHD = congenital heart disease; EPI = epilepsy; ID = intellectual disability. Error bars reflect 95% confidence intervals (Supplementary Appendix G)

### Comparing the *de novo* and epidemiologic effects

Figure 2 presents our estimates of paternal age risk derived through the dnSNV and epidemiologic models, specifically comparing risk for each disorder between fathers older than 39 versus those in their 20s. For the Danish cohort used to estimate the epidemiologic association between advanced paternal age and ASD, SCZ, ID and EPI the mean ages of fathers in their 20s and over 39 were 26.2 and 44.1 respectively. For the cohort used to estimate the epidemiologic association between advanced paternal age and CHD, the mean ages were 27.1 and 43.7 respectively.

In the epidemiologic model, the HR (or OR) for each disorder ranged from: 1.06 (0.89 – 1.18) for CHD to 1.68 (1.51-1.86) for ASD (Table S3). For CHD, EPI and ID the estimates from the epidemiologic and dnSNV models are statistically indistinguishable (p > 0.1 for all comparisons). This means that we do not observe a paternal age effect in the population outside the bounds of what could be explained by paternal-age-related *de novo* variation. For SCZ and ASD, however, the HRs derived from the epidemiologic data were significantly greater than the IRRs expected from paternal-age-related dnSNVs (p =1.4e10^-4^ for SCZ, p < 2e10^-5^ for ASD). This result suggests that the much of the paternal age effect observed in the population is attributable to factors other than *de novo* mutations. The inflated epidemiologic effect was particularly notable for ASDs, as estimated risk in the population was nearly an order of magnitude (9.3X) greater than that which our model could attribute to dnSNVs. Further, the epidemiologic effect observed for ASDs was greater than the epidemiologic effect observed for intellectual disability (p = 0.032). As ID collections have a substantially greater observed rate of *de novo* variants than ASD collections (see Table 1), these data very strongly suggest that factors other than *de novo* variation drive the association between paternal age and autism risk observed in the population.

**Figure 2:**
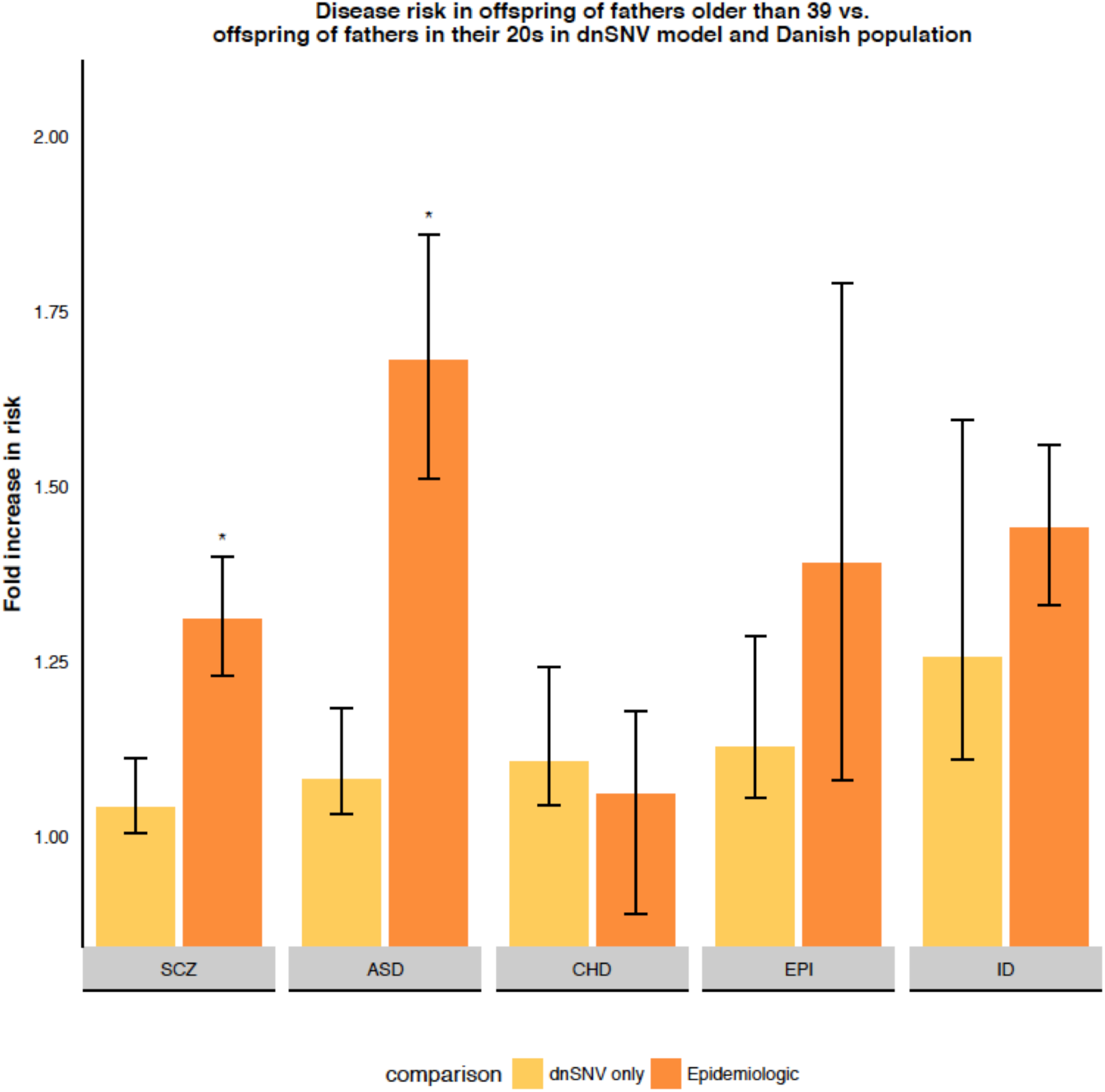
Note: SCZ = schizophrenia; ASD = autism spectrum disorder; CHD = congenital heart disease; EPI = neurodevelopmental disorders with epilepsy; ID = intellectual disability. Error bars reflect 95% confidence intervals. * p < 2*10^-4^ against null hypothesis that epidemiologic association between advanced paternal age and disease risk is equivalent to the dnSNV model’s estimate.

## Discussion

Epidemiologic associations between advanced paternal age and offspring disease-risk are commonly assumed to reflect *de novo* mutations that accumulate inmale gametes as men age. This assumption results in the inference that delaying paternity *causes* these increases in offspring disease risk. In the event that the epidemiologic observations also reflect factors that are time-stable but predict age of childbearing (like inherited genetic liability), the causal interpretation could raise unnecessary concerns among those considering parenthood later in life and be inappropriately troubling to the parents of affected children.

In this analysis, we directly estimated the extent to which paternal-age related *de novo* mutations create offspring disease risk for ASDs, CHD, EPI, ID, and SCZ. For each, we show that the causal effect attributable to *de novo* mutations in the exome is small. By delaying paternity from his mid-20s to his mid-40s, for example, a man’s offspring would be only about 4% more likely to develop schizophrenia, and 29% more likely to develop ID, subsequent to his age-related *de novo* mutations. Against the low prevalence of each condition, this increase in risk results in a small predicted increase in incidence at the population level. Clinicians can use these data to address patient questions about the risks associated with delayed parenting.

For CHD, EPI, and ID, the estimates of risk derived from our dnSNV model were statistically indistinguishable from the epidemiologic estimates from the Danish population analyses. For those conditions, the risk observed in the epidemiologic data is consistent with *de novo*-driven phenomena. This is not true, however, for the behaviorally defined disorders. For ASDs and SCZ, *de novo* mutations do not appear to account for most of the observed epidemiologic patterns.

This finding is consistent with results from a recent simulation study,^15^ as well as research suggesting that a parent’s age at the birth of their first child, rather than the affected child, can explain much of the epidemiologic association between parental age and schizophrenia.^43^ It is also consistent with recent evidence that inherited polygenic risk for disorders such as SCZ and ASD is associated with delayed childbearing.^16,17^

Our estimates could be affected by several additional properties of the data and analysis. First, the trio-sequencing results for four of the disorders (ASD, ID, EPI and CHD) came from datasets that are partially overlapping^18,31^. In actuality, these disorders are highly comorbid as diagnosed, but it is possible that the dnSNV estimates we provide for these disorders are more similar to each other than they would be if totally independent cohorts were used. This limitation cannot be addressed in the present analysis as we employed published summary data from each cohort. Second, while we have explicitly considered the role of *de novo* mutations in the exome, it is possible that *de novo* single nucleotide variants in the rest of the genome could create additional risk associated with advanced paternal age. However, there is currently no evidence for enrichment of *de novo* variants outside of the exome in genetically complex human diseases.^35^ More importantly, there is no reason to expect that such variants, if they did contribute meaningfully, would act only in some disorders (ASD, SCZ) and not others in which the mutational burden in the exome explains the epidemiologic findings (CHD, ID) as we show here.

The average age of childbearing has increased across many communities and cultures. These results should be reassuring in that these trends are unlikely to lead to a substantial increase in common human diseases through the accumulation of *de novo* mutations. In particular, they should lay to rest the belief that an increase in average parental age is a primary driver of the observed increase in incident ASD diagnoses in the United States over the last 30 years.^12^ In fact, our data suggest that an incidence increase of this magnitude would only result from delayed childbearing if American men were, on average, now conceiving their children when well over 100 years old (Supplementary Appendix L).

## Acknowledgements

Thank you to Rosy Hosking for her helpful review. Thank you to Alexander Frieden for his editorial comments. E.B.R. and D.J.W. were funded by National Institute of Mental Health grant 1K01MH099286-01A1 and Brain Behavior Research Foundation (NARSAD) Young Investigator grant 22379. J.L.T. and E.M.W. were funded by the Stanley Center for Psychiatric Research at the Broad Institute. J.P.G.D. was funded by the Danish Council for Independent Research DFF – 1331-00050.H.O.H. was supported by stipends from the Federal Ministry of Education and Research (BMBF), Germany, FKZ: 01EO1501 and the German Research Foundation (DFG): HE7987/1-1. C.E.S. is funded by the Howard Hughes Medical Institute. J.G.S. is funded by the Cardiovascular Development Consortium (1 U01 HL098166). Clinical information and exome data on CHD families was provided by the Pediatric Cardiac Genomics Consortium (U01-HL098188, U01-HL098147, U01-HL098153, U01-605 HL098163, U01-HL098123 and U01-HL098162. We also thank the families who took part in the Pediatric Cardiac Genomics Consortium and the Simons Simplex Collection study and the clinicians who collected data at each of the study sites. The iPSYCH project is funded by the Lundbeck Foundation and the universities and university hospitals of Aarhus and Copenhagen.

